# NheABC is a pH-dependent cytotoxin that contributes to the virulence of *Bacillus cereus*

**DOI:** 10.64898/2026.01.19.700242

**Authors:** Naeem Ullah, Abdelbasset Yabrag, Raswati Pant, Vignesh Ramnath, Toril Lindbäck, Laura M. Carroll, Manoj Puthia, Aftab Nadeem

## Abstract

*Bacillus cereus* is a Gram-positive bacterium widely distributed in food, soil, and plants. It is a spore-forming, facultative anaerobic bacterium known as a food-borne opportunistic human pathogen responsible for causing gastrointestinal and non-gastrointestinal infections, including wound-associated infections. In the present study, we report that among single-component, bi-component, and tripartite alpha pore-forming toxin (α-PFT) producing bacteria, the tripartite NheABC toxin produced by *B. cereus* induced the maximum cell death in infected epithelial cells. Similar to its effects in 2D monolayers, NheABC exhibited potent cell toxicity in 3D spheroids and intestinal organoids, targeting their cell membrane and mitochondria. Moreover, using erythrocytes as a model system, we found that the cytolytic activity of NheABC is pH-dependent, and was markedly reduced at acidic pH (5.5). The pH-dependent biological activity of NheABC was further confirmed by a liposome leakage assay. Importantly, NheABC enhanced the colonization of *B. cereus* in a non-gastrointestinal murine wound infection model. Overall, our study highlights the role of pH in regulating NheABC-mediated cytotoxicity in mammalian cells, which may lead to the development of novel therapeutic strategies for managing *B. cereus* gastrointestinal and non-gastrointestinal infections.

## Introduction

Bacterial infections are a leading cause of morbidity and mortality worldwide. The incidence of lethal bacterial infections, including urinary tract infections, pneumonia, invasive bloodstream infections, skin and soft tissue infections, and wound infections, is rising worldwide ^1, 2^. The pathogenic Gram-positive and Gram-negative bacteria produce pore-forming toxins (PFTs) as a major virulence factor during mammalian host infections that play an important role in facilitating the spread or colonization of pathogenic bacteria within the target host ^2, 3^. Based on the secondary structure of the transmembrane regions responsible for their insertion into lipid membranes, PFTs can be divided into two major groups: α-PFTs (α-helical PFTs) and β-PFTs (β-barrel PFTs). These toxins are secreted in an inactive monomeric form, and upon interaction with the lipid membranes of the target host, they may undergo conformational changes that may trigger their oligomerization resulting in the formation of multimeric pores that are crucial for mammalian cell lysis ^4^. The α-PFTs include the single-component ClyA cytotoxin produced by *Escherichia coli* ^5^, the two-component toxin like XaxAB from *Xenorhabdus nematophila*, XaxAB-like binary toxin from *Photorhobdus luminescens*, the YaxAB from *Yersinia enterocolitica* and the three-component toxins such as MakABE from *Vibrio cholerae*, AhlABC from *Aeromonas hydrophila*, SmhABC from *Serratia marcescens*, Hbl-BL_1_L_2_ and NheABC from *Bacillus cereus* ^6–9^.

*B. cereus* is a Gram-positive, spore-forming, facultative anaerobic bacterium widely distributed in food, soil, and plants. A member of the *B. cereus* group species complex, *B. cereus* is known as a food-borne opportunistic human pathogen responsible for causing gastrointestinal and non-gastrointestinal infections ^10^. The spores of *B. cereus* are highly adhesive and can spread from their natural habitat to food production environments, which may lead to the contamination of any kind of food. The high frequency of food contamination and the production and secretion of the tripartite non-hemolytic enterotoxin (NheABC) may explain the severity of the diarrheal and non-gastrointestinal infections caused by *B. cereus* ^11^. NheABC is a tripartite pore-forming toxin encoded by the genes *nheA, nheB*, and *nheC* ^12^. All the three components of NheA, NheB and NheC in a molar ratio 10:10:1 are required for the full cytolytic activity of the toxin ^13^. The NheABC toxin was first isolated from the culture supernatant of *B. cereus* strain NVH 00075/95, a *panC* Group III/Sequence Type 26 (ST26) strain ^14^ that was responsible for a food poisoning outbreak in Norway in 1995 ^15^ (using *panC* phylogenetic group assignment ^16^ and PubMLST’s seven-gene multi-locus sequence typing scheme for “*B. cereus*”, respectively; note that *B. cereus* group taxonomy is not standardized/debated, but for simplicity, we will refer to NVH 00075/95 as “*B. cereus*”).

For many years, mammalian cell cultures have been extensively used as valuable tools for understanding the complex molecular and cellular mechanisms involved in bacterial infections. The majority of the bacterial infection studies utilize two-dimensional (2D) cell cultures as model systems ^17–19^. The 2D cell culture models have clear limitations, as they do not reflect the complexity of mammalian tissue physiology ^20^. Although the 2D cell culture is inexpensive and highly reproducible, most 2D cell culture systems lack many physiologically relevant features including cell-cell interactions, tight junctions, and the extracellular microenvironment required for the regulation of various cell signalling ^18^. These limitations make the translational value of findings from 2D cell culture less valuable. Therefore, animal models used for understanding the virulence potential of bacterial pathogens may offer a comprehensive understanding of human pathophysiology. Yet, they also come with potential challenges, including high cost, being time consuming, and ethical concerns regarding animal welfare that cannot be ignored. To overcome these short comings, three-dimensional (3D) cell culture models including spheroids and organoids have gained increasing importance in the field of infection biology in recent years ^20–24^. In these models, cells are cultured under *in vitro* conditions that allow them to grow and interact in all three spatial dimensions, closely resembling *in vivo* microstructures. Moreover, some of the cell lines grown as 3D spheroids, and most organoids facilitate the development of cell-extracellular matrix, and cell-cell interactions resulting in tissue architectures that closely resemble *in vivo* conditions, thus making these cellular systems optimal for studying bacterial infections ^23–25^.

In the present study, we report that among the single-component, two-component, and three-component toxin producing bacteria, NheABC produced by *B. cereus* induced the maximum cell death in the infected epithelial cells, which involved targeting the mammalian host cell membrane and mitochondria. Similar to its effects on a 2D monolayer, NheABC caused potent cell toxicity in 3D spheroids and intestinal organoids. Moreover, using erythrocytes as a model system, we found that the biological activity of NheABC is pH-dependent, which was further confirmed by a liposome leakage assay. Importantly, NheABC enhanced the virulence of *B. cereus* in a non-gastrointestinal murine wound infection model.

## Results

### Bacterial pore-forming toxins induce cell death in epithelial cells grown in 2D culture

Numerous pathogenic bacterial species release single-component, two-component, or three-component PFTs. Upon interaction with the host cell membrane, these toxins undergo conformational change, resulting in the formation of an oligomeric pore that leads to cell death^3^. To address the role of these PFTs in the killing of the epithelial cells, we exposed HCT116 cells to various bacterial species at a multiplicity of infection of 50 (MOI: 50) for 4 hours, followed by staining with propidium iodide (dead cells) and Hoechst 33342 (dead and alive cells). Images were acquired using the SparkCyto imaging system. Data analysis suggests that the tripartite, NheABC-producing *B. cereus* induced the maximum cell death in the infected epithelial cells. Like *B. cereus*, *S. marcescens*, another tripartite PFT-producing bacterium, also induced increased epithelial cell death. Importantly, in addition to the tripartite SmhABC, *S. marcescens* is known to produce a β-PFT, ShlA that may also contribute to epithelial cell death ^26^. We used *E. coli* 536, known to produce the β-PFT, hemolysin as a positive control while *B. subtilis*, which lacks PFTs was used as a negative control (**Figure 1A**). The other tripartite (*A. hydrophila*, and *V. cholerae*) and the bipartite PFT-producing bacteria (*Y. enterocolitica*, and *P. luminescens*) failed to induce cell death in the infected epithelial cells (**Figure 1A**).

**Figure 1:**
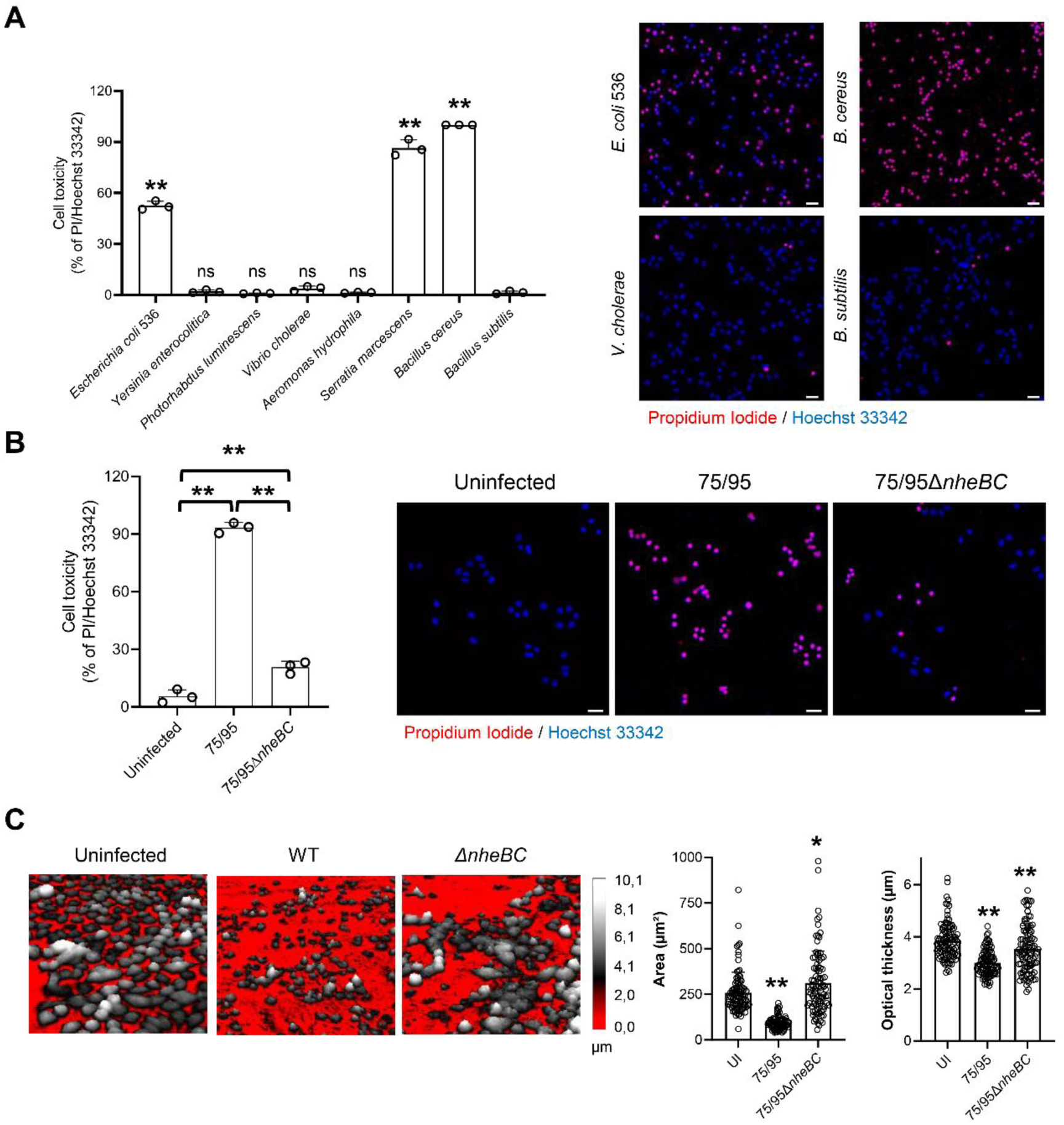
The *B. cereus* NheABC toxin causes potent cell death in epithelial cells. **(A)** HCT116 epithelial cells were infected with various bacterial species (MOI, 1:50) in complete DMEM media for 4 hours at 37°C, followed by staining with propidium iodide (red, staining dead cells), and Hoechst 33342 (blue, staining dead and alive cells). Images were acquired using the SparkCyto imaging system, followed by image segmentation using Cellpose. The total number of propidium iodide-positive dead cells were normalized against the Hoechst 33342-positive total number of cells. Histograms show the percentage of cell death induced by various bacterial species. Data points represent replicates from independent experiments; bar graphs show mean ± s.d. Significance was determined from replicates using a one-way analysis of variance (ANOVA) with Dunnett’s post-test against *B. subtilis*. **p<0.01, *p≤0.05. **(B)** HCT116 cells were infected with wild type *B. cereus* (75/95) and its isogenic mutantΔ*nheBC* (75/95Δ*nheBC*) strain (MOI, 1:50) in DMEM media for 4 hours at 37°C, followed by staining with propidium iodide. Histograms show the percentage of cell death induced by the wild type 75/95 and its isogenic mutant 75/95Δ*nheBC* strain. Data points represent three replicates from independent experiments; bar graphs show mean ± s.d. Significance was determined from replicates using a one-way analysis of variance (ANOVA) with Sidak’s post-test. **p<0.01. **(C)** Changes in HCT8 cell morphology were recorded by holographic microscopy. Cells were infected with wild type 75/95 *B. cereus* and its isogenic mutant 75/95Δ*nheBC* (MOI, 1:50) in DMEM medium for 4 hours at 37°C. The color gradients in the images, from red to white, indicate varying levels of cell thickness, with white representing the thickest cells. The histograms to the right indicate changes in area and optical thickness. Data points in the histogram represent individual HCT8 cells (n=96) from two independent experiments; bar graphs show mean ± s.d. Significance was determined from replicates using a one-way analysis of variance (ANOVA) with Sidak’s post-test against the uninfected cells. **p<0.01, *p≤0.05.

The *B. cereus* NVH75/95 (75/95) is known to produce the tripartite NheABC toxin. In addition, it also has the ability to produce a sphingomyelinase (SMase) that may also contribute to cell death ^27^. To address the role of NheABC in *B. cereus* infection-mediated cell death of epithelial cells, we infected HCT116 cells with the wild type (75/95), and the mutant lacking the NheBC components of the tripartite toxin (75/95Δ*nheBC*). Data analysis suggests that compared to the wild type 75/95 strain, the cytotoxic activity of 75/95Δ*nheBC* was significantly attenuated (**Figure 1B**). Similar results were also observed in the *B. cereus* infected HCT8 cells (**Supplementary Figure 1**). Consistent with the cell death data, we also observed significant changes in the HCT8 cell morphology upon infection with the wild type *B. cereus* 75/95 strain, while its isogenic mutant 75/95Δ*nheBC* strain showed minimal effects (**Figure 1C**). Together these results suggest that NheABC produced by *B. cereus* induces maximum lethality in the infected epithelial cells.

### The NheABC tripartite toxin causes metabolic paralysis of the target epithelial cells

Mitochondrial integrity is essential for cellular homeostasis, and mitochondria have emerged as crucial battlegrounds in the interplay between host defenses and bacterial pathogens ^28^. To investigate whether NheABC produced by *B. cereus* influences the mitochondrial integrity of the epithelial cells, HCT116 cells were infected with the *B. cereus* wild type 75/95, or the isogenic mutant 75/95Δ*nheBC* strain at an MOI of 50 for 4 hours, followed by staining with TMRM, which stains the active mitochondria of mammalian cells (**Figure 2A**). Infection of HCT116 cells with the 75/95 strain showed a significant decrease in TMRM staining compared to the vehicle control. Moreover, HCT116 cells infected with the 75/95Δ*nheBC* strain showed a minimal decrease in the TMRM fluorescence in the infected HCT116 cells. To address whether the loss of mitochondrial potential would lead to a decrease in total cellular ATP, we isolated bacterial-free supernatant from both the wild type 75/95 and the mutant 75/95Δ*nheBC* strain and exposed HCT116 cells to these supernatants. Exposure of HCT116 cells to the supernatant collected from the wild type 75/95 strain caused a time-dependent increase in cell death. Importantly, the cells treated with the supernatants collected from the 75/95Δ*nheBC* strain failed to induce cell death (**Figure 2B**). Consistent with the NheABC-mediated loss of mitochondrial potential and cell death, we observed a time-dependent decrease in the total cellular ATP content of HCT116 cells exposed to the wild-type 75/95 compared to the 75/95Δ*nheBC* strain (**Figure 2C**). To determine whether loss of mitochondrial function leads to morphological changes in epithelial cells, we treated HCT8 cells with bacteria-free supernatant from the wild-type 75/95 strain. This treatment induced mitochondrial rounding, demonstrated by a reduction in mitochondrial length compared to vehicle-treated cells (**Figure 2D-E**). Together, these results suggest that the NheABC tripartite toxin targets both the cell membrane and mitochondria of the target epithelial cells.

**Figure 2:**
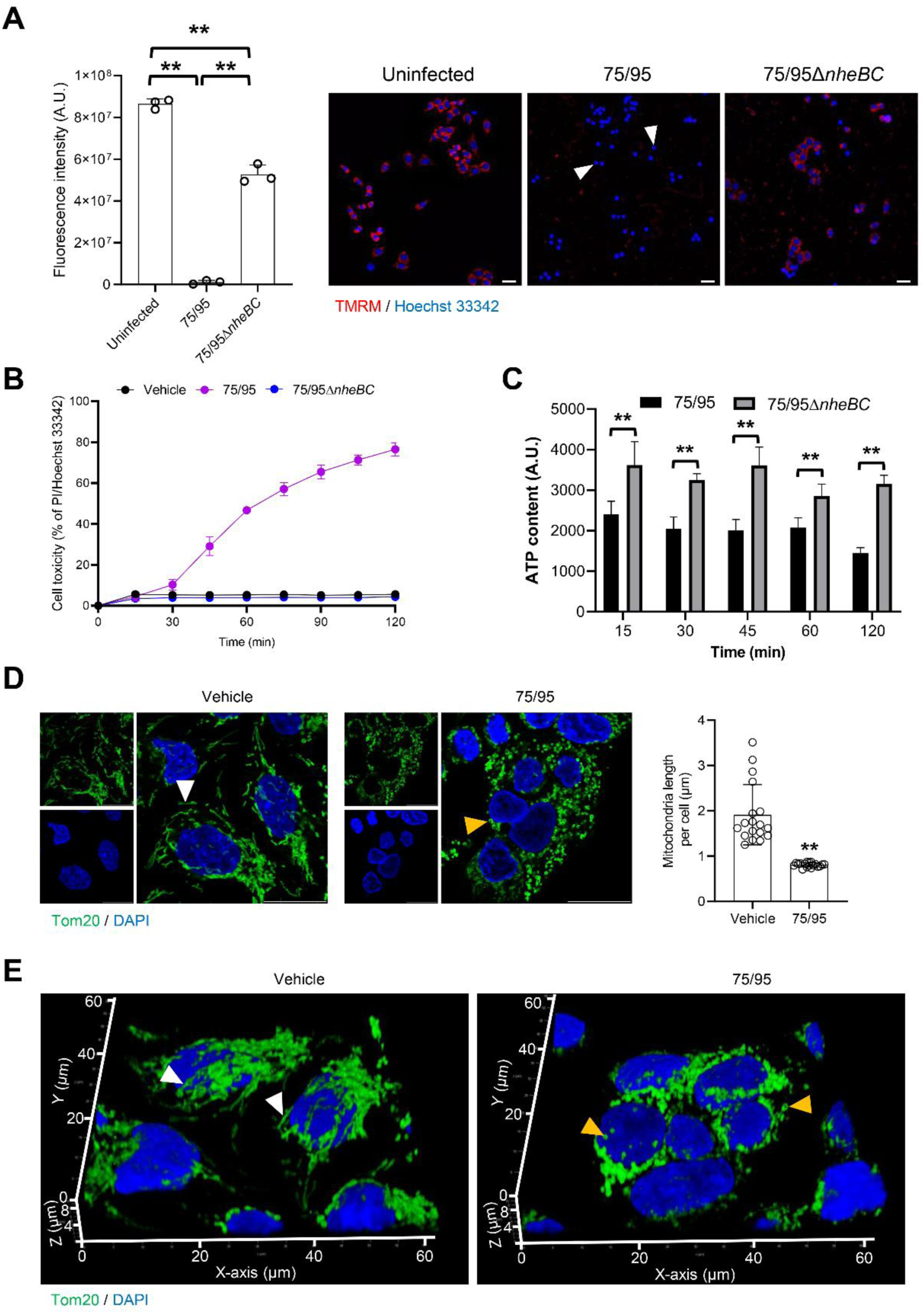
The tripartite toxin NheABC contributes to the metabolic paralysis of epithelial cells. **(A)** HCT116 cells were infected with wild type *B. cereus* (75/95) and its isogenic mutant Δ*nheBC* (75/95Δ*nheBC*) strain (MOI, 1:50) in DMEM medium for 4 hours at 37°C, followed by staining with TMRM (red), and Hoechst 33342 (blue). Arrowheads (white) indicate a loss of mitochondrial potential. Data points in the histogram to the left represent fluorescence intensity of TMRM staining (1650 x 1650 µm^2^) from independent experiments; bar graphs show mean ± s.d. Significance was determined from replicates using a one-way analysis of variance (ANOVA) with Sidak’s post-test against the uninfected cells. **p<0.01. **(B)** DLD1 cells pre-stained with Hoechst 33342 (2 µM) in RPMI-1640 media were exposed to bacteria-free supernatants (5 %) collected from the wild type *B. cereus* (75/95) and its isogenic mutant Δ*nheBC* (75/95Δ*nheBC*) strain. For vehicle control, cells were treated with LB (5 %). Images were acquired every 15 min after treatment, and the number of propidium iodide positive cells was counted as dead cells. **(C)** DLD1 cells were exposed to bacteria-free supernatants (5 %) collected from the wild type *B. cereus* (75/95) strain or its isogenic mutant Δ*nheBC* (75/95Δ*nheBC*). Cells were lysed in a 96-well plate every 15 min, for a maximum of 120 min to calculate total cellular ATP. Data points represent three replicates from independent experiments; bar graphs show mean ± s.d. Significance was determined from replicates using unpaired Student’s t-test. **p<0.01. **(D-E)** HCT8 cells were exposed to vehicle (5% LB) or bacteria-free supernatant (5%) from the wild type *B. cereus* (75/95) for 30 min. Mitochondria were detected with anti-Tom20 antibody (green), and nuclei were counterstained with DAPI (blue). White arrowheads indicate healthy tubular mitochondria, while orange arrowheads indicate round, fragmented mitochondria. The histogram to the right in panel (D) shows a decrease in mitochondria length in response to supernatant from wild-type *B. cereus*. Data points represent quantification of mitochondrial length from 18 random cells; bar graphs show mean ± s.d. Significance was determined using unpaired Student’s t-test. **p<0.01. The images in panel (E) show representative 3D reconstructions of mitochondria in HCT8 cells treated with vehicle or wild-type *B. cereus* supernatants.

### NheABC produced by *B. cereus* causes cell death in 3D spheroids and organoids

Two-dimensional (2D) cell culture models have been extensively used as a tool to understand the cellular mechanisms involved in *B. cereus* PFT-mediated mammalian cell death ^13^. These models have a clear limitation as they lack the complex cell-cell interactions, and the extracellular microenvironment ^18, 24^. Therefore, the use of three-dimensional (3D) cell culture models, including spheroids and organoids may offer an alternative to the simplistic 2D cell culture methods. To investigate whether *B. cereus* producing the NheABC toxin may cause cell death in the infected 3D spheroids, we first developed spheroids from HCT116 cells and infected them with the wild type *B. cereus* 75/95, or the 75/95Δ*nheBC* strain for 4 hours, followed by propidium iodide and Hoechst 33342 staining (**Figure 3A and Supplementary Figure 2**). The wild type *B. cereus* 75/95 strain caused disruption of the HCT116 cell spheroid, as indicated by an almost complete loss of the spheroid core. Moreover, we also observed an increase in the number of propidium iodide positive cells in the *B. cereus* 75/95 infected HCT116 spheroids, compared to the uninfected spheroid (**Figure 3A**). To further understand the kinetics of NheABC tripartite-mediated cell killing in the HCT116 spheroids, we infected the spheroids with either the wild type *B. cereus* 75/95, or 75/95Δ*nheBC* strain in the presence of propidium iodide. Images were acquired using the SparkCyto imaging system. Data analysis suggested a time-dependent increase in propidium iodide staining for the HCT116 cells exposed to the *B. cereus* 75/95 strain, compared to the 75/95Δ*nheBC* infected spheroids (**Figure 3B**). To further address if NheABC may cause disruption of the 3D spheroid, we infected the HCT116 spheroids with *B. cereus* 75/95, followed by staining with propidium iodide and Hoechst 33342. Consistent with the SparkCyto image analysis, confocal microscopy analysis suggested an increase in the number of propidium iodide-positive cells in the spheroids infected with *B. cereus* 75/95. Importantly, we also observed an increase in the FITC-Dextran permeability of the spheroid infected with the wild type *B. cereus* 75/95, compared to the uninfected spheroid (**Figure 3C**). Together, these results suggest that NheABC is the major virulence factor responsible for the disruption of HCT116 spheroids during *B. cereus* infection.

**Figure 3:**
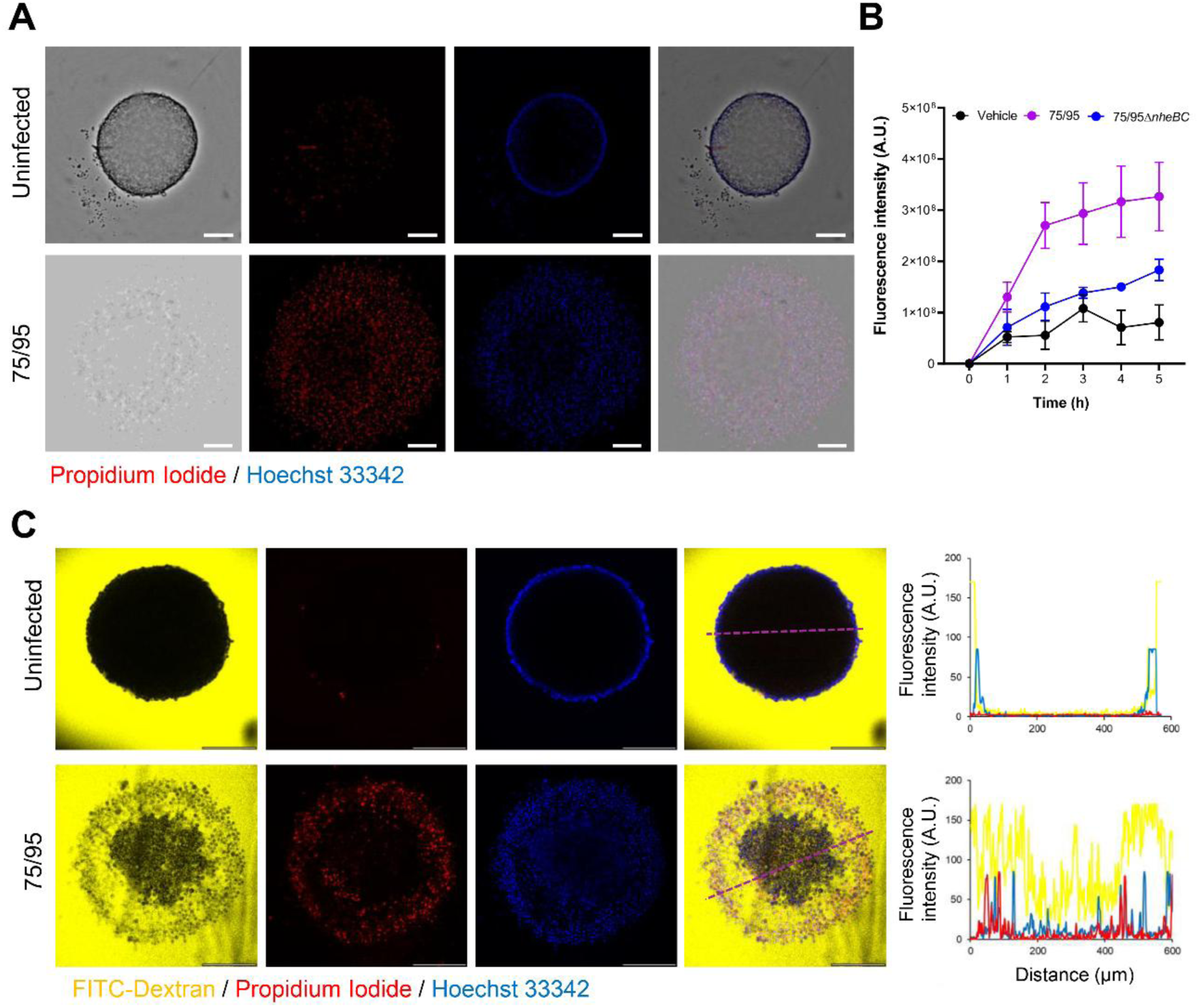
*Bacillus cereus* causes disintegration of spheroids prepared from HCT116 cells. **(A)** Spheroids formed from HCT116 cells for 72 hours in a 96-well round bottom plate, followed by their infection with wild type *B. cereus* (75/95) for 4 hours in an antibiotic-free medium at 37°C. At the end of infection spheroids were stained with propidium iodide (red), and Hoechst 33342 (blue). **(B)** The line graph shows a time-dependent increase in the propidium iodide fluorescence in spheroids infected with the wild type *B. cereus* (75/95) strain, or its isogenic mutant 75/95Δ*nheBC*. The line graph shows data from three independent spheroids; line plot shows mean ± s.d. **(C)** The HCT116 spheroids formed in a 96-well plate were transferred to an 8-well glass bottom slide and infected with the wild type *B. cereus* (75/95) for 4 hours. The spheroids were stained with propidium iodide (red), Hoechst 33342 (blue), and FITC-Dextran (yellow). The line plot to the right indicates an increase in the number of propidium iodide positive cells and an increase in the penetration of FITC-Dextran into the lumen of the infected spheroid, compared to the uninfected spheroid.

We next investigated the role of NheABC-mediated cell death in the intestinal organoids, which are composed of various types of differentiated cells that self-assemble into a small, organ-like structures ^29^. The intestinal organoids were infected with the wild type *B. cereus* 75/95, or 75/95Δ*nheBC* strain for 4 hours, followed by staining with propidium iodide and Hoechst 33342. Images were acquired using the SparkCyto imaging system. Data analysis suggests that organoids infected with *B. cereus* 75/95 showed an increase in cell death compared to the organoids infected with 75/95Δ*nheBC* (**Figure 4A-B**). To determine whether *B. cereus* may cause mitochondrial dysfunction in intestinal organoids, similar to that in the cells grown in 2D monolayers, we infected organoids with *B. cereus* 75/95, followed by staining with TMRM and Hoechst 33342. Images were acquired using confocal microscopy suggested a marked decrease in TMRM staining in organoids infected with *B. cereus* 75/95, compared to the uninfected organoids. Taken together, the data clearly demonstrate that *B. cereus* 75/95 causes depolarization of mitochondria in the infected organoids (**Figure 4C-D**). These results suggest that, like in 2D cell culture, the *B. cereus* NheABC toxin targets the cell membrane and mitochondria of the target cells in the spheroids and organoids.

**Figure 4:**
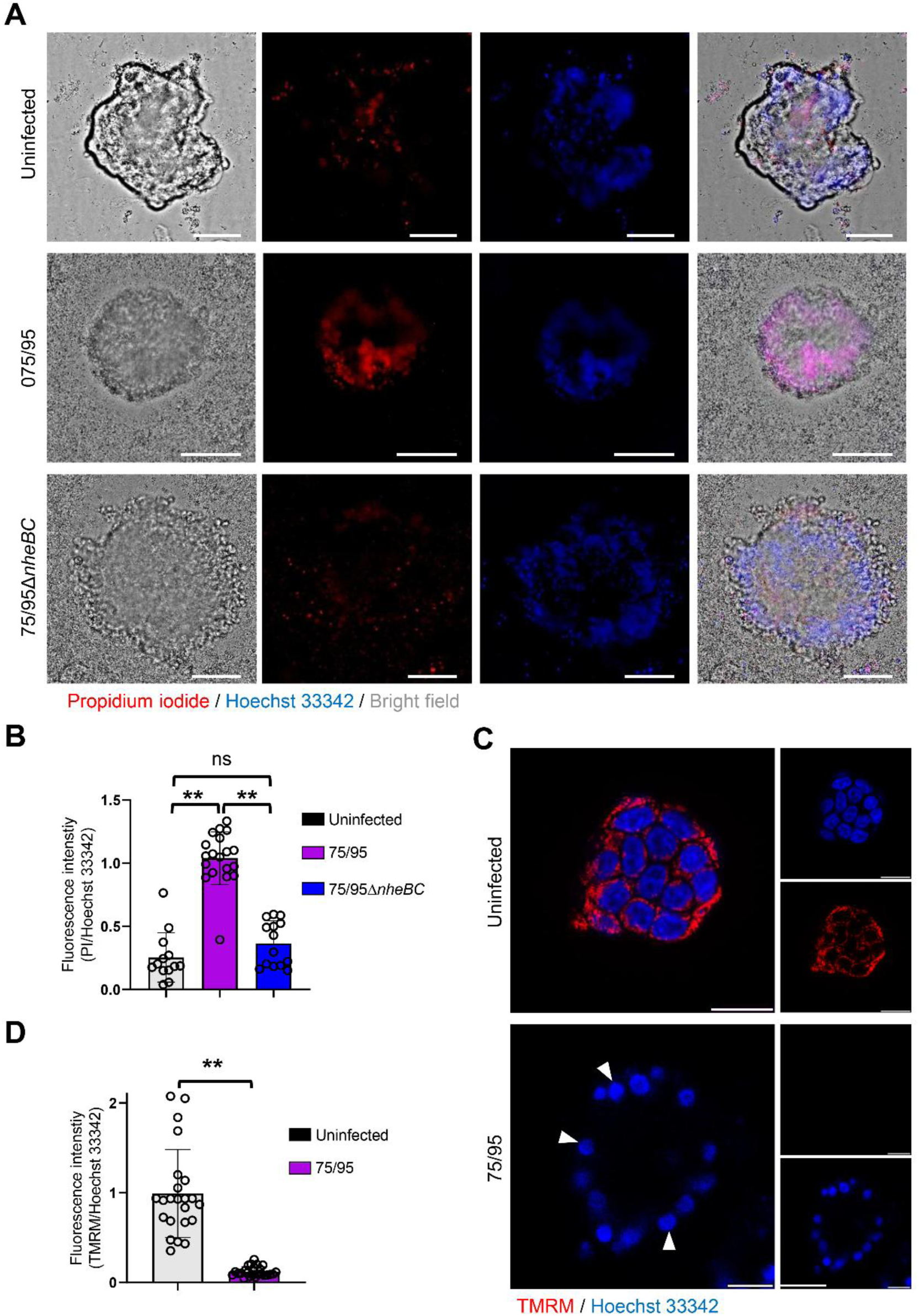
*Bacillus cereus* targets the cell membrane and mitochondria of the infected intestinal organoid. **(A)** Intestinal organoids were infected with the wild type *B. cereus* (75/95), or its isogenic mutant 75/95Δ*nheBC* strain for 4 hours, followed by staining with propidium iodide (red), and Hoechst 33342 (blue). **(B)** Histograms indicate an increase in the fluorescence intensity of propidium iodide in organoids infected with the wild type *B. cereus* (75/95), compared to its isogenic mutant 75/95Δ*nheBC* strain or uninfected organoids. Data points represent quantification of fluorescence from individual intestinal organoid (n = 13 to 19); bar graphs show mean ± s.d. Significance was determined using a one-way analysis of variance (ANOVA) with Sidak’s post-test against the uninfected cells. **p<0.01, ns = non-significant. **(C)** Intestinal organoids were infected with the wild type *B. cereus* (75/95) for 4 hours, followed by staining with TMRM (red), and Hoechst 33342 (blue). Arrowheads (white) indicate loss of mitochondrial potential. **(D)** Histograms indicate a decrease in the fluorescence intensity of TMRM staining in organoids infected with the wild type *B. cereus* (75/95), compared to the uninfected organoids. Data points represent quantification of individual intestinal organoid (n = 23); bar graphs show mean ± s.d. Significance was determined using Student’s t-test. **p<0.01.

### Targeting of the mammalian cell membrane by NheABC is pH-dependent

The NVH75/95 strain of *B. cereus* is known to produce SMase in addition to NheABC as an additional virulence factor ^27^. To address if both NheABC and SMase produced by *B. cereus* are active against mammalian cells, we collected bacteria-free supernatants from *B. cereus* 75/95 and 75/95Δ*nheBC*, following their exposure to human erythrocytes for 4 hours. Data analysis from erythrocytes hemolysis suggests that *B. cereus* 75/95 caused almost complete hemolysis, unlike 75/95Δ*nheBC* which remained inactive in this assay (**Figure 5A**). We then investigated the kinetics of human erythrocyte cell lysis using a turbidity assay (**Figure 5B, and Supplementary Figure 3A-B**). Consistent with the hemolytic assay, we observed a concentration– and time-dependent decrease in erythrocyte turbidity (indicative of an increase in hemolysis) in response to the supernatants collected from the *B. cereus* 75/95 strain, while supernatants from the 75/95Δ*nheBC* strain remained inactive (**Figure 5B and Supplementary Figure 3A-B**). Taken together, the results suggest that NheABC is the primary virulence factor responsible for the lysis of human erythrocytes.

**Figure 5:**
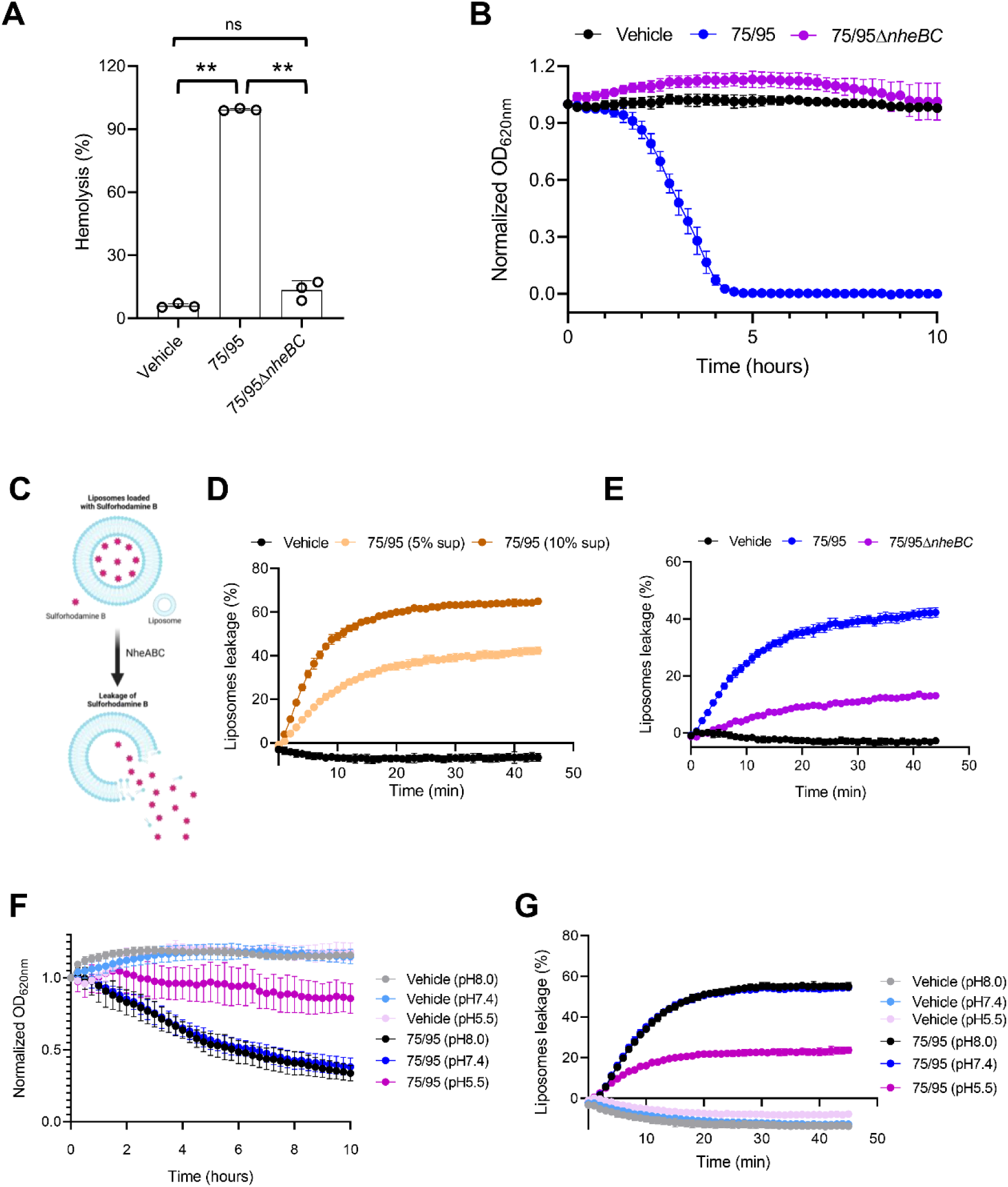
NheABC causes pH-dependent lysis of the target cell membrane. **(A)** Human erythrocytes were exposed to bacteria-free supernatants collected from the wild type *B. cereus* (75/95) and its isogenic mutant Δ*nheBC* (75/95Δ*nheBC*) strain for 4 hours at 37°C, followed by quantification of hemolysis at 415 nm. Histograms show the percentage of hemolysis induced by the wild type 75/95 and its isogenic mutant 75/95Δ*nheBC* strain. Data points represent three replicates from independent experiments; bar graphs show mean ± s.d. Significance was determined using a one-way analysis of variance (ANOVA) with Sidak’s post-test against the vehicle treated cells. **(B)** A turbidity assay was used to determine the kinetics of erythrocytes lysis by measuring its optical density every 10 minutes after the addition of the bacteria-free supernatants (5% v/v) collected from the wild type strain, *B. cereus* (75/95) or its isogenic mutant *ΔnheBC* (75/95Δ*nheBC*) for 6 hours at 37°C. LB (5% v/v) was used as a vehicle control. **(C)** Schematic illustration of a liposome leakage assay performed with liposomes loaded with sulforhodamine B prepared from the epithelial cell lipid extract (ECLE). **(D)** The line plot indicates concentration– and time-dependent ECLE liposome leakage induced by the wild-type *B. cereus* (75/95) supernatants in citrate buffer (120 mM, pH7.4). **(E)** The line plot indicates time-dependent ECLE liposome leakage induced by the supernatants isolated from wild-type *B. cereus* (75/95) or its isogenic mutant Δ*nheBC* (75/95Δ*nheBC*) strain in citrate buffer (120 mM, pH7.4). **(F)** A turbidity assay was used to determine the kinetics of erythrocytes lysis by measuring the optical density of human erythrocytes every 10 minutes after the addition of the bacteria-free supernatants (5% v/v) collected from the wild type *B. cereus* (75/95) or its isogenic mutant Δ*nheBC* (75/95Δ*nheBC*) strain for 10 hours at 37°C. LB (5% v/v) was used as a vehicle control. Buffer refers to citrate buffer (120 mM) of the indicated pH. **(G)** The line plot indicates time-dependent ECLE liposome leakage induced by the wild-type *B. cereus* (75/95) supernatant in citrate buffer (120 mM), at the indicated pH. Data in panels D-G are representative of two or more independent experiments. Data presented in the graphs is representative of three or more replicates.

To determine the role of mammalian cell lipids in NheABC-mediated mammalian cell lysis, we extracted lipids from the epithelial cell lipid extracts (ECLE) and loaded them with sulforhodamine B (**Figure 5C**). We exposed sulforhodamine B loaded liposomes to increasing concentration of bacteria-free supernatants. Data analysis suggests a concentration-dependent increase in the leakage of liposomes prepared from the ECLE (**Figure 5D**). To determine if SMase present in the bacteria-free supernatant may also cause leakage of the ECEL liposomes, we exposed the liposomes to bacteria-free supernatants collected from *B. cereus* 75/95 or the 75/95Δ*nheBC*. Data analysis suggests that unlike human erythrocytes, the liposomes prepared from ECLE were sensitive to the SMase produced by *B. cereus* 75/95Δ*nheBC* (**Figure 5E**). Taken together, our results suggests that both human erythrocytes and ECLE liposomes are sensitive to the NheABC tripartite toxin. The partial leakage of ECLE liposomes in response to 75/95Δ*nheBC* could be explained due to the potential presence of SMase in the bacteria-free supernatants.

Upon ingestion of contaminated food, *B. cereus* colonizes the small intestine of the infected individuals ^30^. Thus, the ability of the *B. cereus* to adapt to a slightly acidified environment is crucial to its virulence against the mammalian host. The pH of the small intestine and colon may reach as low as 6.0 ^31^. To investigate the role of pH in NheABC-mediated lysis of erythrocytes, we exposed erythrocytes to various pH conditions (8.0 to 5.5) and quantified erythrocyte lysis by a turbidity assay. Data analysis suggests that erythrocyte lysis was maximum at pH 8.0 and reduced under acidic pH conditions (**Figure 5F**). Consistent with the decrease in NheABC-mediated lysis of human erythrocytes, we also observed a reduction in NheABC-mediated leakage of liposomes prepared from the ECLE (**Figure 5G**). Taken together, the data suggests that acidic pH reduces NheABC-mediated hemolytic activity against erythrocytes and leakage of ECLE liposomes.

### Diverse *B. cereus* group strains have been isolated from wounds in humans

Previous studies have reported that the pH of a wound infections ranges between 7.15 and 8.9 ^32^. *B. cereus* is a well-described causative agent of food poisoning. In addition, *B. cereus* has long been associated with traumatic wound infections ^33–36^. However, the role of NheABC in *B. cereus* wound infection is not well known. Using BTyperDB (a *B. cereus* group genomic database with manually curated metadata) ^37^, we identified 41 genomes of good-to-high quality, which were reportedly isolated from wounds in humans (**Table S2**). Notably, these 41 genomes were diverse and spanned multiple species (**Figure 6A**). Specifically, using *panC* Groups as a proxy for species ^16^, the wound-associated strains belonged to *panC* Groups IV (18 genomes, 43.9%), III (11 genomes, 26.8%), II (9 genomes, 22.0%), and V (1 genome; 2.4%); 2 genomes (4.9%) could not be assigned to a *panC* Group with confidence but were placed among genomes from *panC* Groups II and III using whole-genome-based methods (i.e., Mashtree and BTyperDB’s average nucleotide identity [ANI]-based methods; **Figure 6A**). BTyper3 (as implemented in BTyperDB) was used to identify genes encoding known *B. cereus* group toxins within each of the 41 wound-associated genomes (**Table S2**). Using default amino acid identity and coverage thresholds (i.e., 70% and 80%, respectively), none of the wound-associated genomes (0%) possessed anthrax toxin-encoding *cya*, *lef*, and *pagA*, nor did any possess cereulide synthetase-encoding genes *cesABCD* (**Table S2**). As observed previously for the entirety of the *B. cereus* group ^30, 38^, nearly all genomes possessed Nhe-encoding *nheABC* (40/41, 97.6%; **Figure 6A**). Interestingly, one *panC* Group IV genome (i.e., strain SJ-S28, isolated in Memphis, Tennessee, United States, from a human skin infection) ^39^ lacked all of *nheABC* at default amino acid identity and coverage thresholds (BTyperDB ID BTDB_2022-0002555.2, NCBI BioSample accession SAMN07351956; **Figure 6A**); even when both thresholds were lowered to 0%, *nheABC* could not be detected (**Table S3**).

**Figure 6:**
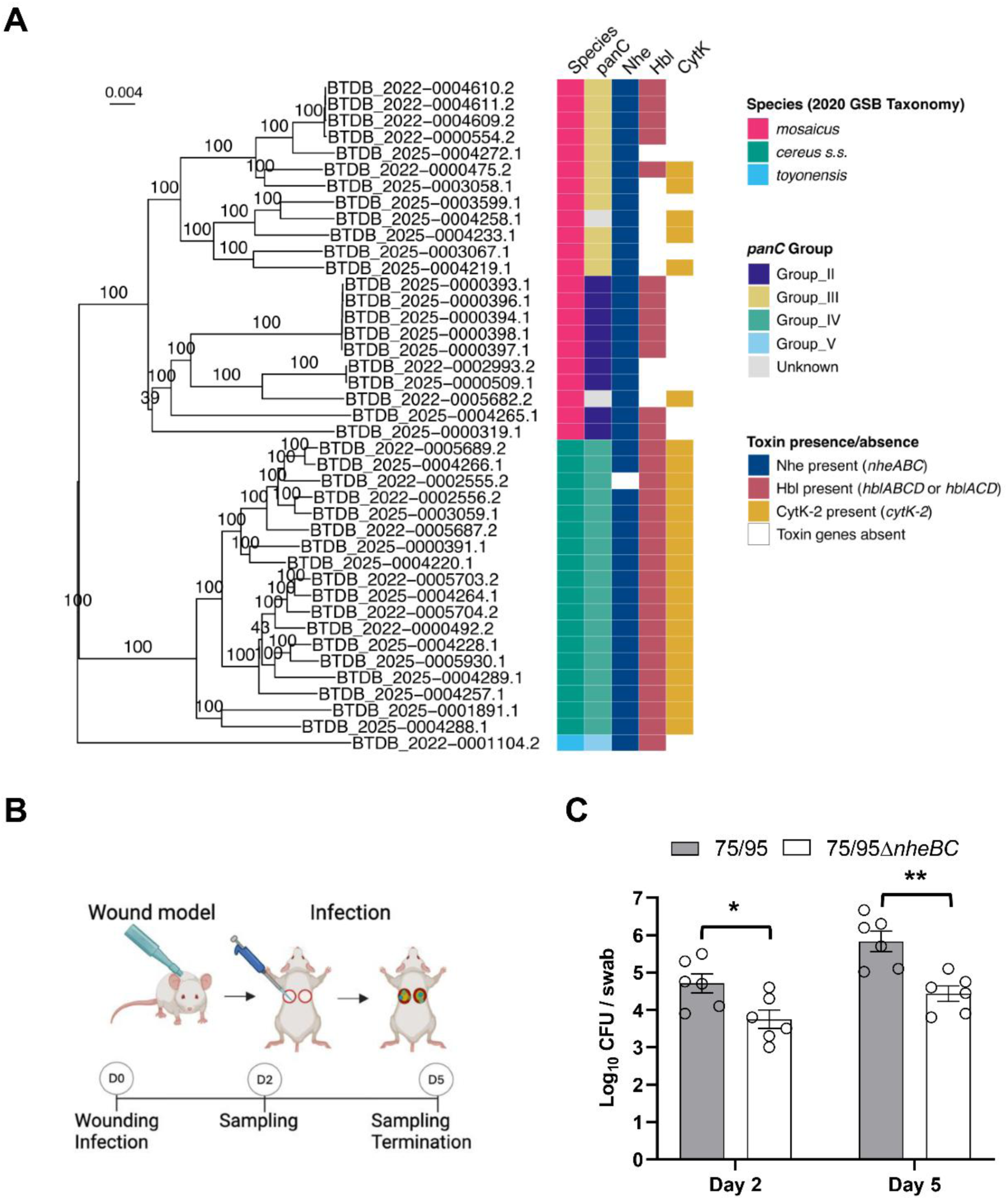
NheABC contributes to virulence of *B. cereus* in wound infections. **(A)** Diverse *B. cereus* group strains have been isolated from wounds in humans. The distance-based tree (left) was constructed using Mashtree and displays 41 *B. cereus* group genomes, which were reportedly isolated from wounds/lesions in humans. The heatmap (right) denotes (from left to right): (i) *B. cereus* group genomospecies (obtained using BTyper3’s 2020 Genomospecies-Subspecies-Biovar [GSB] Taxonomy); (ii) *panC* Group (obtained using BTyper3); the presence and absence of genes encoding (iii) non-hemolytic toxin (Nhe; *nheABC*), (iv) hemolysin BL (Hbl; either *hblABCD* or *hblACD*), and (v) cytotoxin K variant 2 (CytK-2; *cytK-2*), all detected using BTyper3 and default thresholds (i.e., 70% and 80% amino acid identity and coverage, respectively). The Mashtree distance-based tree is rooted at the midpoint, with branch labels denoting bootstrap support percentages (out of 100 repetitions). Bootstrap support percentages on some internal nodes have been omitted for readability (see Supplementary Figure 4 for the raw, unedited version of this figure). **(B)** Schematic illustration of the wound infection experiment. **(C)** Histogram indicates recovery of wild-type *B. cereus* (75/95) or its isogenic mutant Δ*nheBC* (75/95Δ*nheBC*) 2– and 5-days post-infection from the wound of the infected mice. Data points in the histogram represents CFU counts for bacteria collected from the individual mouse; bar graphs show mean ± s.d. Significance was determined using unpaired Student’s t-test. *p<0.01, **p<0.01.

In addition to Nhe, genes encoding Hbl (*hblABCD* and *hblACD*) were detected in the majority (31, 75.6%) of the 41 wound-associated *B. cereus* group genomes (**Figure 6A**). Specifically, all genomes from *panC* Groups IV and V possessed genes encoding Hbl, whereas Hbl-encoding genes were variably present within *panC* Group II and III genomes (similar to results observed previously for *panC* Groups II-V) ^38^. Variant 2 of the gene encoding CytK (*cytK-2*) was additionally detected in a majority of the genomes queried here (24 of 41 wound-associated genomes, 58.5%; **Figure 6A**). Like Hbl (and similar to results observed previously) ^38^, CytK-2-encoding *cytK-2* was present in all *panC* Group IV strains and variably present within *panC* Groups II and III; however, *cytK-2* could not be detected in the wound-associated *panC* Group V genome at default thresholds (**Figure 6A**).

### NheABC contributes to successful colonization of *B. cereus* in wound infections

Wound infections caused by *B. cereus* remain a public health concern ^40^. To date, little is known about the role of NheABC in non-gastrointestinal wound infections. To determine the role of NheABC in wound infections, we used a mouse model of excisional wound and infected the wounds with either the wild type *B. cereus* 75/95 or 75/95Δ*nheBC* strain (**Figure 6B**). The CFU count from the infected wounds suggests a time-dependent increase in bacterial load in mice infected with *B. cereus* 75/95 compared to those infected with 75/95Δ*nheBC*. The wild type *B. cereus* 75/95 showed an increase in colonization at both 2 days and 5 days post-infection, compared to the 75/95Δ*nheBC* strain (**Figure 6C**). Taken together, the results suggest that, similar to gastrointestinal infections, NheABC plays an important role in non-gastrointestinal wound infections.

## Discussion

Pathogenic bacteria produce PFTs as major virulence factors against mammalian cells. In the present study, we report that among the single-component, bi-component, and tripartite PFTs-producing bacteria, NheABC produced by *B. cereus* induced the maximum cell death in the infected epithelial cells, which involved targeting of the mammalian host cell membrane and mitochondria. Similar to its effects on 2D monolayer, the NheABC caused potent cytotoxicity in 3D spheroids and intestinal organoids. Moreover, using erythrocytes as a model system, we found that the biological activity of NheABC is pH dependent. The pH-dependent biological activity of NheABC was further confirmed by a liposome leakage assay. Importantly, our bioinformatics analysis suggests that *B. cereus* isolates from human wound infections had a high frequency of NheABC (97.6 %). Moreover, the presence of the NheABC toxin enhanced the virulence of *B. cereus* in a murine wound infection model.

During the last decade, 3D cell culture has rapidly advanced our understanding of tumor and infection biology, showing promising outcomes ^24^. The 3D spheroids and organoid models closely mimic animal and human tissue, thereby aiding in the reducting animal model usage. In the current study, we used 3D spheroids prepared from HCT116 monolayers and intestinal organoids as a model system to investigate the role of NheABC-mediated targeting of mammalian cell membranes and mitochondria. The cellular mechanisms involved in *B. cereus*-mediated mitochondrial damage, however, need further investigation. We sought to determine if spheroids and organoids could be used as an alternative to the 2D cell culture model to understand the cellular mechanisms involved in PFT-mediated cellular responses in complex cellular systems that closely mimic *in vivo* conditions. Our results suggest that similar to 2D cell cultures, NheABC targets mammalian cell membranes and mitochondria in complex organoid cell cultures. Moreover, we were able to translate these findings to an *in vivo* murine wound infection model, where we observed NheABC-mediated enhanced colonization of *B. cereus* in the infected wounds. These results suggest that NheABC may have a potential role in *B. cereus* invasion into deep tissues.

In previous studies, several PFTs have been reported to have an increased acidic-pH mediated interaction with lipid membranes, such as the MakA cytotoxin ^7, 41, 42^ from *V. cholerae*, a component of the MakABE cytotoxin, colicin A from *E. coli* ^43^, listeriolysin O (LLO) from *Listeria monocytogenes* ^44^ vacuolating cytotoxin A (VacA) from *Helicobacter pylori* ^45^, and perfringolysin O (PFO) from *Clostridium perfringens* ^46^. Most of these PFTs undergo a pre-pore to pore transition under low pH conditions, thereby activating their cytolytic activities. Moreover, the effect of pH on the pore-forming ability of two *B. thuringiensis* toxins, Cry1Ac and Cry1C has been previously reported ^47^. Cry1Ac is highly effective at alkaline pH, while Cry1C is active in an acidic to neutral pH envrionment ^47^. The putative pH-sensing mechanism in NheABC is still unclear, but it likely relates to the protonation of specific amino acids, such as histidines at low pH ^41^. Importantly, it has previously been reported that in addition to α-PFTs such as MakA, a member of β-PFTs LLO is activated by acidic pH (<6). At neutral pH, both MakA and LLO exhibit very little cytotoxic activity, but their activity is substantially increased at low pH (5.5). In contrast, NheABC activity substantially decreased at low pH (5.5), while maximum cell toxicity was observed at pH 7.4 and above. A plausible explanation for the pH-dependent decrease in NheABC activity could be that acidic pH may inhibit the association of the NheABC tripartite components with the lipid membranes. However, this needs further investigation.

In the human gastrointestinal tract, pH is a key factor in shaping gut microbial composition and activity. The pH increases substantially from the stomach (pH1.5 to 3.5) to the small intestine (pH5.5 to 8.0), providing more favorable conditions for most of the microorganisms, including the pathogenic bacterium, *B. cereus* ^48^. To our knowledge, the importance of pH on the biochemical properties of *B. cereus*’s major virulence factor, the NheABC toxin, has not been previously investigated. Our cellular and biochemical data strongly suggest that the cytotoxic activity of NheABC is high at slightly alkaline pH, while a drop in pH towards acidic pH significantly decreases its cytolytic activity. These findings suggest that NheABC may act as a major virulence factor in bacterial infection sites where the pH is slightly alkaline. Previous studies have reported that the pH of a wound infections ranges between 7.15 and 8.9 ^32^. *B. cereus* is a well-described causative agent of food poisoning. In addition, there have been many reports of non-gastrointestinal infections caused by *B. cereus* in humans and animals. It has long been associated with traumatic wound infections ^33–36^. However, the role of NheABC in *B. cereus* virulence in wound infection is not well known. The present study investigates the role of NheABC in *B. cereus* pathogenesis, demonstrating its toxicity in 2D and 3D cell cultures. The pH-mediated activity against mammalian cell membranes suggests that NheABC favours neutral to alkaline environments typical of wound infection sites. By using advanced *in vitro* cellular models with *in vivo* validation, we provide evidence that NheABC not only targets mammalian cell membranes and mitochondria but also enhances the colonization of *B. cereus* in murine wound infections. These findings suggest that targeting the pH-mediated mechanisms of NheABC may lead to the development of novel therapeutics for treating gastrointestinal and non-gastrointestinal infections caused by *B. cereus*.

## Material and Methods

### Growth conditions for bacterial strains

All bacterial strains used in this study are listed in **Table S1**. The bacterial strains were grown on Luria/Lysogeny agar (LA) plates and incubated overnight at 37°C.

### Cell cultures

HCT116, HCT8, and DLD1 cell lines were maintained in RPMI-1640 media (Sigma-Aldrich) supplemented with 10% fetal bovine serum (FBS), 1% penicillin/streptomycin, and non-essential amino acids at 37°C with 5% CO_2_.

### Ethics statement

All animal experiments were performed according to Swedish Animal Welfare Act SFS 1988:534 and were approved by the Animal Ethics Committee of Malmö/Lund, Sweden. The study was conducted in accordance with the local legislation and institutional requirements.

### Bacterial infections and cell toxicity

The bacterial strains were grown on Luria Agar (LA) plates at 37°C overnight, as specified above. The HCT116, and HCT8 cell lines were grown in RPMI-1640 medium supplemented with 10% fetal bovine serum (FBS) and incubated at 37°C overnight in an incubator with continuous flow of 5% CO_2_. The cells were grown in a 96-well plate (1×10^4^ cells/well), or 24-well plate (5×10^4^ cells/well). The following day, cells were washed with PBS and infected with the indicated bacteria (MOI, 1:50) for 4 hours in a serum free DMEM or RPMI-1640 media. Cell toxicity was measured by staining the cells with a mixture of propidium iodide (1 µg/mL) and Hoechst 33342 (2 µM) for 30 min, followed by image acquisition using the SparkCyto imaging system (Tecan), and analysis using ImageJ ^49^ or Cellpose ^50^. A customized python code was developed for batch processing of images acquired with SparkCyto imaging system. The Cyto2 segmentation model ^50^ was used for these analyses to ensure accurate and efficient segmentation of the cell populations.

### Kinetics of epithelial cell death and estimation of total cellular ATP

DLD1 cells (1×10^4^ cells/well) were grown in a 96-well plate and incubated at 37°C and 5% CO_2_ overnight. The following day, DLD1 cells were incubated with a mixture of propidium iodide (1 µg/mL) and Hoechst 33342 (2 µM) for 30 min and exposed to bacteria-free supernatants (5 %) collected from the wild type *B. cereus* 75/95 strain or its isogenic mutant 75/95 Δ*nheBC*. Images were acquired on SparkCyto imaging system every 15 min for a maximum duration of 120 min at 37°C with a continuous flow of 5% CO_2_. The acquired images were analysed as discussed above.

Total cellular ATP contents were measured in DLD1 cells. Briefly, DLD1 cells (1×10^4^ cells/well) were grown in a 96-well plate and incubated at 37°C and 5% CO_2_ overnight. The following day, DLD1 cells were exposed to bacteria-free supernatants (5 %) collected from the wild type *B. cereus* 75/95 strain or its isogenic mutant 75/95 Δ*nheBC*. Cells were lysed in a 96-well plate every 15 min, for a maximum duration of 120 min, to calculate total cellular ATP. Cellular ATP levels were quantified using the ATPLite kit (PerkinElmer, #6016943).

### *B. cereus* infection and mitochondrial potential

HCT116 cells were grown in a 96-well plate (1×10^4^ cells/well) in RPMI-1640 medium supplemented with 10% fetal bovine serum (FBS) and incubated at 37°C overnight in an incubator with continuous flow of 5% CO_2_. The following day, cells were washed with PBS and infected with wild type *B. cereus* 75/95 and its isogenic 75/95Δ*nheBC* mutant strain (MOI, 1:50) for 4 hours in a serum free DMEM media. At the end of infection cells were stained with mitochondrial potential dye TMRM (100 nM) and Hoechst 33342 (2 µM) for 30 min, followed by image acquisition using the SparkCyto imaging system (Tecan), and analysis using ImageJ ^49^.

### Preparation of 3D spheroids and bacterial infection

To develop spheroids, HCT116 (ATCC) cells were grown in a cell repellent 96-well plate (BIOFLOA 96-well cell culture plate [#F202003]) for 72 hours using Cancer Stem Premium (#20141-500) supplemented with non-essential amino acids. The spheroids were infected with the wild type *B. cereus* 75/95 and its isogenic 75/95Δ*nheBC* mutant strain (10^4^ bacteria) for 4 h at 37°C. Cell toxicity was measured by staining the cells with a mixture of propidium iodide (1 µg/mL) and Hoechst 33342 (2 µM) for 30 min, followed by image acquisition using the SparkCyto imaging system (Tecan). Images acquired with the SparkCyto imaging system and analyzed using ImageJ ^49^.

### Preparation of intestinal organoids culture and bacterial infection

Colon intestinal organoids were purchased from MEMD Millipore, an affiliate of Merck KGaA (14-881CR) SCC310. Matrigel matrix basement membrane (Corning, #356234) was used for the preparation of the dome. Human basal medium (IntestiCult OGM, #100-0150) and organoids supplement (#100-0151) were purchased from Stem Cell Technology and used as medium for the maintenance of organoid culture. Gentle cell dissociation reagent (#100-0485, Stem Cell Technology organoids) was used to dissociate organoids from the dome. Organoids (*n* = 20–30) were infected with the wild type *B. cereus* 75/95 and its isogenic mutant Δ*nheBC* (75/95Δ*nheBC*) strain (10^4^ bacteria) in Cancer Stem media supplemented with 1% Matrigel for 4 hours at 37°C. Cell toxicity was measured by staining the intestinal organoids with a mixture of PI (1 µg/mL) and Hoechst 33342 (2 µM) for 30 min, followed by image acquisition using the SparkCyto imaging system (Tecan). Images were analyzed using ImageJ ^49^.

### Holographic microscopy

Holographic microscopy was performed using the HoloMonitor® M4 (Phase Holographic Imaging AB) equipped with a motorized stage to investigate the changes in epithelial cell morphology. HCT8 cells were cultured overnight in RPMI-1640 medium in a 96-well (1×10^4^) or 24-well (5×10^4^ cells/well) plates. The following day, cells were infected with wild type *B. cereus* 75/95 and its isogenic 75/95Δ*nheBC* mutant strain (MOI, 1:50) for 4 hours at 37°C, after which images were acquired using the HoloMonitor® M4. This system generates label-free images reconstructed into three-dimensional holograms. Quantitative measurements, such as average cell thickness and area, were extracted using Hstudio™ software ^51^.

### Confocal microscopy

For the preparation of cells for confocal microscopy, HCT116 spheroids and intestinal organoids were infected with the wild type *B. cereus* 75/95 or its isogenic mutant Δ*nheBC* (75/95Δ*nheBC*) strain (10^4^ bacteria) for 4 hours in an 8-well chamber slide with a coverslip bottom (μ-Slide, ibidi). HCT116 spheroids were stained with a mixture of propidium iodide (1 µg/mL), Hoechst 33342 (2 µM), and FITC-Dextran 70kDa (0.2 mg/mL) for 30 min. Intestinal organoids were stained with TMRM (100 nM), and Hoechst 33342 (2 µM) for 30 min.

Organoids were infected with the wild-type *B. cereus* 75/95 (10^4^ bacteria) in Cancer Stem media supplemented with 1% Matrigel for 4 hours at 37°C. Loss of mitochondrial potential was investigated by staining the organoids with TMRM (100 nM), and Hoechst 33342 (2 µM) for 30 min, followed by image acquisition using the confocal microscopy.

To investigate mitochondrial morphology, HCT8 cells (1X10⁴ cells/well) were grown overnight on an 18-well, glass-bottom chamber slide (μ-Slide, ibidi) at 37°C. The following day, the cells were exposed for 1 hour to either vehicle (5% LB) or 5% bacteria-free supernatant collected from the wild-type B. cereus strain 75/95. Subsequently, the cells were fixed with paraformaldehyde and permeabilized with 0.25% Triton X-100 in PBS. They were then incubated overnight at 4°C with Tom20 antibodies (BD Biosciences, #612278; 1:100 in 5% FCS-PBS) and washed three times with PBS. After washing, the cells were incubated with Alexa-488-conjugated secondary antibodies (Thermos Fischer, 1:200) for 1 hour at room temperature. Nuclei were counterstained with DAPI (Sigma Aldrich, 1 µg/mL 5% FCS-PBS) for 5 minutes and washed three times with PBS. Images were acquired using a Leica SP8 inverted confocal system (Leica Microsystems).

Fluorescence intensity profiles were generated using the plot profile function in ImageJ, and image processing was performed using the ImageJ–FIJI distribution ^49^.

### Human erythrocytes lysis assay

For kinetic measurements of erythrocyte lysis, human erythrocytes (0.25% v/v) were exposed to bacteria free supernatants in a phosphate buffered saline (PBS) at indicated concentrations. Erythrocytes were treated with 5% (v/v) of bacteria free supernatant from either wild-type *B. cereus* 75/95 or the isogenic mutant, 75/95Δ*nheBC* strain. Erythrocytes treated with 5% (v/v) LB was used as a negative control, while erythrocytes treated with TritonX-100 (0.025% v/v) were used as positive control. Turbidity of the erythrocytes was monitored every 15 minutes by measuring optical density at 620 nm using Spark microplate reader (Tecan) with intermittent orbital shaking at 37°C.

The kinetics of erythrocytes cell lysis under various pH condition (pH 8.0, pH 7.4, and pH 5.5) in response to bacteria free supernatant from wild-type *B. cereus* 75/95 was performed in various pH-adjusted citrate buffers (120 mM).

The percentage (%) of erythrocyte cell lysis was calculated using the following formula: The diluted erythrocytes were added into the 96-well microplate wells, and turbidity of the samples was measured as F_0_. After the addition of 5 μl of bacteria free supernatant, the turbidity was continuously measured as *F_tn_* every 15 min for 10 hours. The lowest turbidity after the addition of TritonX-100 was measured as *F_100_*. The erythrocytes lysis is defined in two steps as, Step1: F_1_ = (F0 or *F_tn_*–*F_100_*), and Step2: F*_tn_*/F*_1_*. Each erythrocytes lysis assay was performed in three to four technical replicates. The experiment was repeated at least three times.

### Liposomes extractions and leakage assay

Epithelial cell lipid were extracted from three 75 cm² confluent flasks of DLD1 colon cancer cells using the Folch method ^52^. Briefly, cells were harvested, pelleted, and resuspended in a 2:1 chloroform-methanol mixture to extract lipids. The organic phase was collected, and the lipid extract was dried under a gentle nitrogen stream to form a thin lipid film, yielding approximately 5 mg of dried lipids.

To investigate leakage of liposomes prepared from epithelial cell lipid extract (ECLE) in response to NheABC, sulforhodamine B (SRB) leakage assay was performed. SRB was dissolved at 50 mM in a sodium citrate buffer (120 mM). Lipid films (1 mg) were rehydrated with 1 mL SRB solution. The hydrated lipid suspensions were extruded 11 times through a 100 nm polycarbonate membrane using an Avanti Mini-Extruder (Avanti Polar Lipids) at 40°C to form unilamellar liposomes. Unbound SRB-dye from the SRB loaded liposomes was removed using Sephadex G-50 chromatography.

To perform liposome leakage assay in response to bacteria-free supernatants collected from wild-type *B. cereus* 75/95 or its isogenic mutant 75/95Δ*nheBC* strain, SRB-encapsulated liposomes were diluted in 20 times with citrate buffer (120 mM, pH 7.4). The kinetics of liposomes leakage was investigated using Spark plate reader (Tecan).

The kinetics of ECLE leakage assay under various pH condition (pH 8.0, pH 7.4, and pH 5.5) in response to bacteria free supernatant from wild-type *B. cereus* 75/95 was performed. The fluorescence of sulforhodamine B acid was measured in a Spark plate reader with excitation of 540 nm and emission filter of 620 nm. The 20 times diluted liposomes were added into the 96-well microplate wells, and the fluorescence emission were recorded as *F_0_*, 5 μl of bacteria free supernatant was added into each well, and the fluorescence emission was continuously measured as *F_tn_* every 1 min for 45 to 60 min. At the end of the experiment, 5 μl of 0.25% TritonX-100 was added to each well for maximum leakage of the liposomes. Addition of 5 μl of LB was used as a negative control. The highest fluorescence emission after the addition of TritonX-100 was measured as *F_100_*. The percentage of liposome leakage is defined by the following formula, [(Ftn – F0) / F100] × 100. Each liposome leakage assay was performed in three to four technical replicates. The experiment was repeated at least three times.

### Acquisition of human wound-associated *Bacillus cereus* group genomes

Human wound-associated *B. cereus* group genomes and their associated metadata were acquired from BTyperDB ^37^. Briefly, all genomes meeting the following criteria were downloaded from BTyperDB (version 2025 November 30): (i) the strain was isolated from a human (‘Source_1 == “Human”’); (ii) the strain was responsible for an “Other” type of illness (i.e., not foodborne and not anthrax; ‘Human_Illness == “Other”’) or an unknown type of illness (‘Human_Illness == “Unknown”’); (iii) the strain’s metadata contained one of more of the following key words, in any field (case insensitive): ‘wound’, ‘lesion’, ‘skin infection’; (iv) the genome was not classified by BTyperDB as “low quality” (‘Genome_Quality != “Low_Quality“’); (v) the genome did not belong to *B. anthracis* (i.e., *B. mosaicus* subsp. *anthracis*, using the 2020 Genomospecies-Subspecies-Biovar [GSB] taxonomy ^53^ assigned via BTyperDB’s implementation of BTyper3 ^38^; to ensure strains from cutaneous anthrax cases were removed, ‘BTyper3_subspecies != anthracis’). The metadata for genomes that resulted from this search was manually inspected to confirm that each strain was isolated from a human wound. Overall, this search resulted in 41 human wound-associated genomes, which were used in subsequent steps (**Table S2**).

### Tree construction and visualization

Mashtree v1.4.5 ^54^ was used to construct a distance-based tree using the 41 human wound-associated *B. cereus* group genomes as input (see section “Acquisition of human wound-associated *Bacillus cereus* group genomes” above) and the following parameters: (i) the ‘mashtree_bootstrap.pl’ command; (ii) 100 bootstrap repetitions (‘--reps 100’); (iii) the ‘--mindepth’ parameter set to 0 (to prioritize tree accuracy by ignoring very unique *k*-mers that are more likely read errors; ‘--mindepth 0’); (iv) a genome size of 5,619,812 bp (i.e., the median size of the 41 human wound-associated genomes; ‘--genomesize 5619812’); (v) 8 CPUs (‘--numcpus 8’).

The resulting tree was plotted in R v4.4.0. Specifically, the Newick file produced by Mashtree was loaded into R using the ‘read.treè function from ape v5.8-1 ^55^ and rooted at the midpoint using the ‘midpoint.root’ function from phytools v2.4-4 ^56^. Metadata from BTyperDB was loaded into R using the ‘read.delim’ function (parameters: ‘header = T, sep = “\t”, stringsAsFactors = F, check.names = F’). The midpoint-rooted tree was plotted using the ‘ggtreè function from ggtree v3.12.0 ^57^. The ‘gheatmap’ function from ggtree was used to display the following metadata from BTyperDB in a heatmap next to the tree: (i) species (per BTyper3’s 2020 GSB Taxonomy); (ii) *panC* Group (per BTyper3); (iii) presence/absence of genes encoding Nhe (*nheABC*), Hbl (*hblABCD* or *hblACD*), and CytK (*cytK-1* or *cytK-2*; detected using BTyperDB’s implementation of BTyper3, default settings).

### Toxin gene detection in an Nhe-negative genome

According to BTyperDB, one genome (strain SJ-S28, BTyperDB ID BTDB_2022-0002555.2, NCBI BioSample accession SAMN07351956) lacked Nhe-encoding *nheABC* (a nearly ubiquitous toxin within the *B. cereus* group) ^30, 38^. To confirm the absence of *nheABC*, the SJ-S28 genome was queried for toxin genes separately, using BTyper3 v3.4.0 ^38^ and amino acid identity and coverage thresholds lowered to 0 (‘--virulence_identity 0 –-virulence_coverage 0’). The resulting tblastn table was manually inspected to confirm that *nheABC* were absent (**Table S3**).

### Bacterial culture preparation for wound infection

Both wild type *B. cereus* 75/95 and 75/95Δ*nheBC* strains were used for wound infection experiments. A single bacterial colony was inoculated in a tube with 5 mL of Todd Hewitt Broth (THB) and incubated overnight at 37 °C in a shaking incubator. To refresh the culture the next day, 100 μL of the overnight culture was inoculated into a tube containing 5 mL of THB. The tube was then incubated at 37 °C in a shaking incubator until the optical density (OD) reached 0.4. Cultures were then centrifuged at 5,600 rpm for 10 min. The bacterial pellet was washed in 5 mL Tris buffer (10 mM, pH 7.4) and centrifuged again. The resulting pellet was then diluted in Tris buffer to a density of 10⁸ CFU/mL and used to infect wounds.

### Mouse model of wound infection

To study wound infection, a mouse model of excisional wound was used. BALB/c mice (8–10 weeks old male; Janvier Labs, France) were anesthetized by intraperitoneal injection of ketamine (60 mg/kg) and xylazine (10 mg/kg). The dorsal skin was shaved with clippers, and a depilatory cream was applied to remove any remaining hair. The skin was then disinfected with 10% povidone-iodine solution followed by wiping with a 70% alcohol swab.

A sterile 4 mm biopsy punch was used to create markings for two wounds—one on the left and one on the right side of the midline—with at least 5–6 mm distance between the two. Circular skin pieces were then excised with scissors. Wounds were covered with sterile gauze pieces. Thirty minutes later, mice were given 1 ml of prewarmed saline subcutaneously. For analgesia, buprenorphine (0.1 mg/kg mixed in saline) was administered.

Wounds were infected with 10 µL of bacterial suspension (10^8^ CFU/mL). The wounds were then covered with a polyurethane dressing (Mepilex Transfer; Mölnlycke, Sweden), and a transparent adhesive film dressing (Tegaderm; 3M) was applied.

Under isoflurane anesthesia, observations were performed, and wound dressings were changed every other day. The experiment was terminated on day 5. During observation, sterile cotton swab samples were collected from wounds for further microbiological analysis. Swab bacterial CFU analysis was done as described before ^58^.

## Statistical analysis

Data are shown as mean ± s.d. Statistical significance was determined by one-way ANOVA, Student’s t-test or otherwise stated in the corresponding figure legends. Statistical significance is indicated as p ≤ 0.01 (**) or p ≤ 0.05 (*). Graphs were produced and statistical analysis were performed using GraphPad Prism software.

## Acknowledgements

A.N. received support from the Swedish Research Council (2022-04779) and the Kempe Foundations (JCSMK23-0138). V.R. and L.M.C. were supported by the SciLifeLab & Wallenberg Data Driven Life Science (DDLS) Program (grant: KAW 2020.0239) and the Swedish Research Council (grant: 2023-05212). M.P. acknowledges grants from Edvard Welanders stiftelse och Finsenstiftelsen (Hudfonden), and Alfred Österlunds stiftelse. We acknowledge the facilities and technical assistance of the Umeå Core Facility Electron Microscopy (UCEM) and the Biochemical Imaging Center (BICU), Umeå University, a part of the National Microscopy Infrastructure NMI (VR-RFI 201600968 and VR-RFI 2019–00217). We acknowledge the Protein Expression and Purification facility (PEP) at Umeå University for construct design and cloning. Genomic computation was enabled by resources provided by the National Academic Infrastructure for Supercomputing in Sweden (NAISS), partially funded by the Swedish Research Council through grant agreement no. 2022-06725, as well as High Performance Computing Center North (HPC2N; Umeå University, Umeå, Sweden).

## Author Contributions

A.N. conceived the project and wrote the initial draft of the manuscript. All co-authors contributed to the design of experiments, data analysis, interpretation, manuscript revisions, and agree on the final contents of the manuscript.

## Author Information

Correspondence and requests should be addressed to A.N.

