## Supplementary Information for "NheABC is a pH-dependent cytotoxin that contributes to the virulence of *Bacillus cereus*"

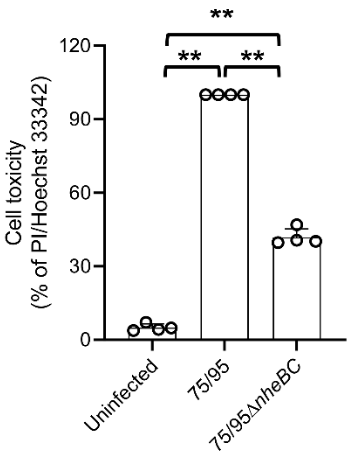

**Supplementary Figure 1: NheABC contributes to epithelial cell death in response to *B. cereus* infection**

HCT8 cells were infected with wild type *B. cereus* (75/95) and its isogenic mutant *ΔnheBC* (75/95*ΔnheBC*) strain (MOI, 1:50) in DMEM medium for 4 hours at 37°C, followed by staining with propidium iodide. Histograms show the percentage of cell death induced by the wild type 75/95 and its isogenic mutant 75/95*ΔnheBC* strain. Data in the histogram is representative of two biological replicates. Data points represent data from four technical replicates; bar graphs show mean ± s.d. Significance was determined from replicates using a one-way analysis of variance (ANOVA) with Sidak's post-test. **p<0.01.

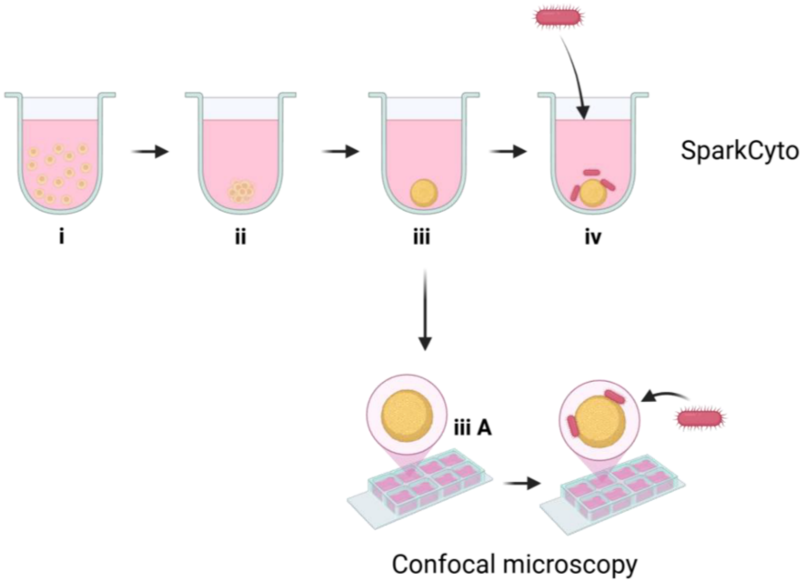

**Supplementary Figure 2: Schematic illustration of steps involved in spheroids formation, and *B. cereus* infection**

**(i)** HCT116 cells were added to a 96-well plate. **(ii)** 12-24 hours after addition to the 96-well plate, HCT116 cells form a loose aggregate. **(iii)** 72 hours after seeding HCT116 cells, a compact spheroid was formed that were ready for infection experiment. **(iii-A)** The compact spheroids formed in a 96-well plate were transferred to an 8-well cover-slip bottom chamber slide, followed by infection with B. cereus for 4 hours and confocal microscopy. **(iv)** Infection of HCT116 spheroids with *B. cereus* in a 96-well plate.

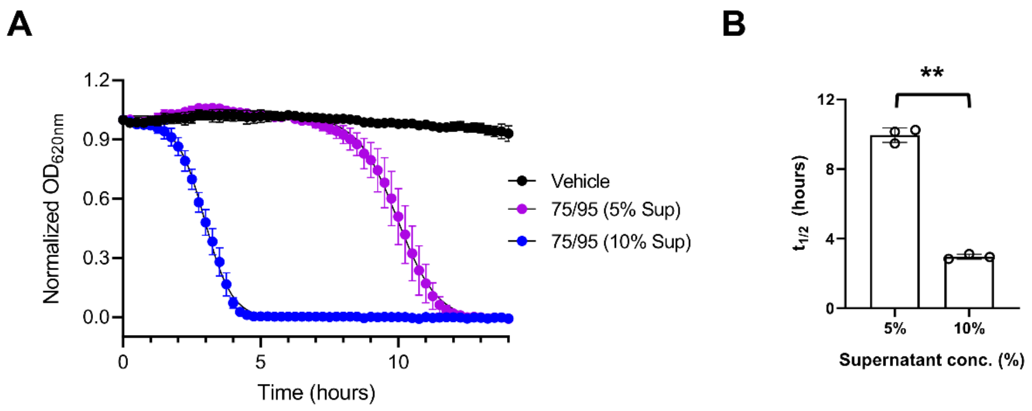

**Supplementary Figure 3: *B. cereus* supernatant induces dose-dependent lysis of human erythrocytes.**

**(A)** A turbidity assay was used to determine the kinetics of erythrocytes lysis by measuring the optical density of human erythrocytes every 10 minutes after the addition of increasing concentration of bacteria-free supernatants collected from the wild type *B. cereus* (75/95) for 14 hours at 37°C. LB (10% v/v) was used as vehicle control. **(B)** The $t_{1/2}$ of decrease in erythrocytes turbidity was reduced by approximately three-fold when erythrocytes were exposed to 5% bacteria-free supernatant compared to the 10% bacteria-free supernatant.

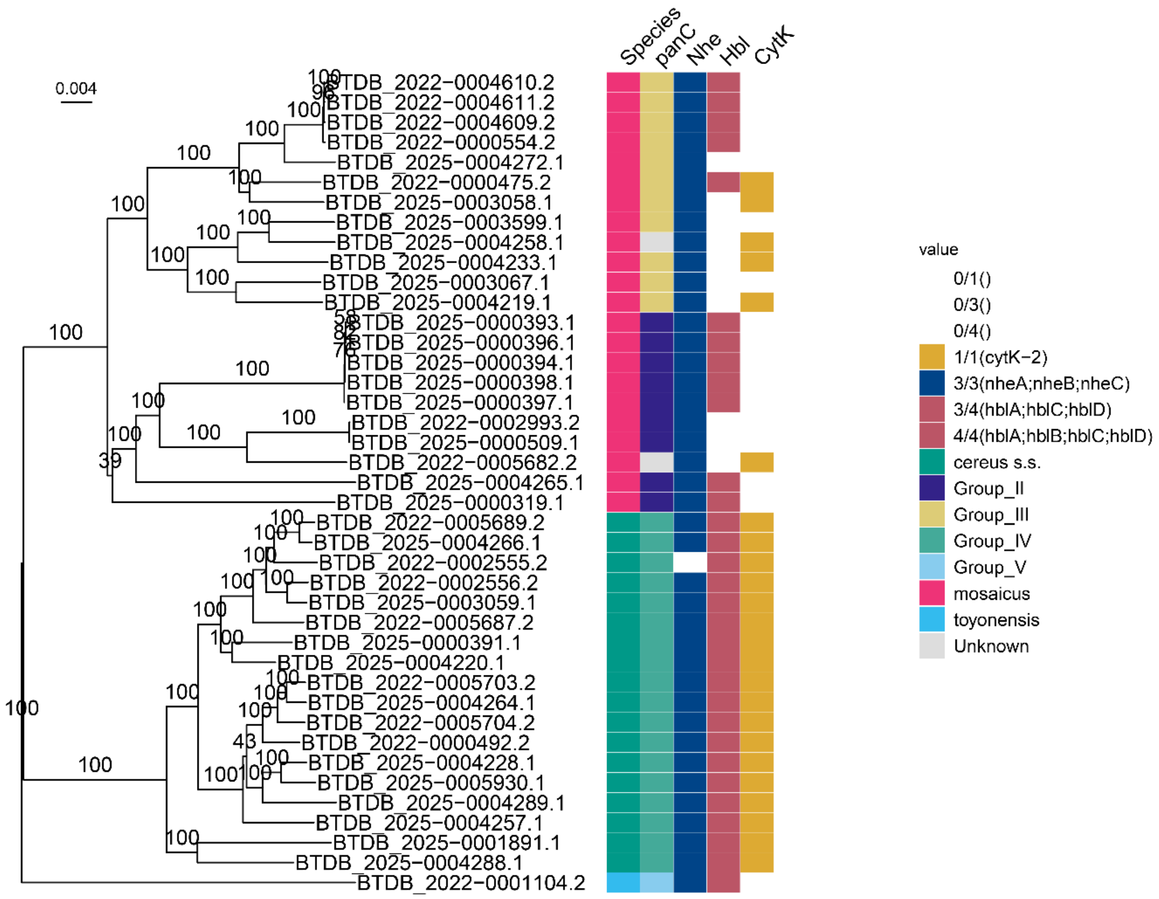

**Supplementary Figure 4: Unedited version of figure 6A**

**Table S1: Bacterial species used in the study**

| Strains | Description/Relevant characteristics | Reference |
|---|---|---|
| *Escherichia coli* 536 | UPEC strain (O6:K15:H31) wild type | [1] |
| *Yersinia enterocolitica* | Wild type | W22703 |
| *Photorhabdus luminescens* | Wild type | TT01 |
| *Serratia marcescens* | Wild type | ATCC 274 |
| *Vibrio cholerae* A1552 | Serogroup O1, $Rif^R$ | [2] |
| *Aeromonas hydrophila* AH3 | Serotype O:34 | [3] |
| *Bacillus cereus* | Wild type (NVH0075/95) | [4] |
| *Bacillus cereus ΔnheBC* mutant | *ΔnheBC* (NVH0075/95) | [5] |
| *Bacillus thuringiensis* BT 407 | Wild type | [6] |
| *Bacillus subtilis* 168 | BGSC, Bacillus Genetic Stock Centre, Columbus | [7] |
